# Paleochronic reversion in *Psophocarpus*, the decompression function in floral anatomic fields

**DOI:** 10.1101/070540

**Authors:** Edward. G.F. Benya

**Author notes:** http://www.egfbenya.com ORCHID iD: 0000-0002-8566-8346 doi: http://dx.doi.org/10.1101/070540.

## Abstract

Paleochronic reversion (an atavism) in *Psophocarpus* presents a basic floral phylloid ground state. That ground state can quickly change as permutation transformation (T_x_) begins. The form of permutation can vary as phyllotactic phylloid (T_Phyld_) and/or floral axial decompression (T_Axl_) presenting linear elongation (T_Long_), rotational (T_Rtn_) and/or lateral (T_Lat_) components. Research with 70 reverted floral specimens documented varying degrees of phyllotactic permutation at the bracts (Bt) region and inter-bracts (IBS) sub-region of the pre-whorls pedicel-bracts anatomic zone. Permutation further yielded an inter-zonal pericladial stalk (PCL). It continued at the floral whorls zone: the calyx (Cl), corolla (Crla), androecium (Andr), and gynoecium (Gynec) with components therein. These organ regions present a continuum as an axial dynamic vector space £Taxi of floral permutation dominated by axial expansion (AE) so that an anatomic sequence of permutation activity runs from the bracts (Bt) region to the carpel (Crpl) inclusive with components therein, summarized by the formula:

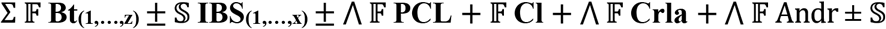 stamen fltn 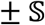 Andr spiral 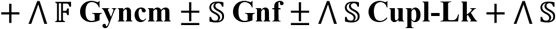 Crpl ±_(Crpl web ± VASCARP ± Crpl diadn ± Crpl fltn + [fltn no] ± Crpl Rtn)_ = T_x_. The flower reverts from a system of determinate growth to one of indeterminate growth.

## 1. Introduction

Shoot apical meristem (SAM) genesis follows a biophysical compressed, cylindrical form whose structuring function is highly specific (Besnard et al. 2014). Classic studies have documented a “steady-state approximation for cylindrical shoot models” (Young, 1978) of the SAM whose single organ constitution, phyllotaxis and development (i.e. leaf) presents a specificity of form, at minimal variation, that is captured and summarized with precision in a single complete model (Green and Baxter, 1987; Young, 1978). The resulting compact, organ sequence of leaves presents biophysical fields as nodes and internodes whose identity is verified at bud-burst and bloom. Permutative internodal elongation (T_Long_) decompression reveals phyllotactic order that, although variable, is precise (Jeune and Barabé, 2006).

That structure and exactitude change as the SAM undergoes “evocation or induction” (transformation “T_x_”) to a floral meristem (FM) whose organ composition amplifies from a single leaf morphology in the SAM to multiple organ morphologic forms of converted leaves (Battey and Lyndon, 1990; Ditta, et al. 2004; Surridge, 2004; Weigel and Meyerowitz, 1994). In most angiosperms (Stern, 1988) those converted leaves appear at two specific floral anatomic zones of pre-whorl (i.e. pedicel and bracts) and of whorls (i.e. calyx, corolla, androecium and gynoecium). The FM maintains a compact, compressed cylindrical sequence of organs, similar to the SAM, but whose exactitude of form and sequence can be affected by homeotic genes (Coen and Meyerowitz, 1991; Goto et al. 2001; Honma and Goto, 2001; Ikeda et al. 2005; Kidner and Martienssen, Li et al. 2017; 2005; Pautot et al. 2001). The FM thus presents a structure similar to that of the SAM but as a reproductive system (Stern, 1988) whose organs usually distribute in biophysical fields of specific spirals and/or whorls regions (Endress and Doyle, 2007). Thus precision of the FM is less than that of the SAM. However it is still significantly specific and is summarized by a dynamic; the ABC(DE) model (Coen and Meyerowitz, 1991; Honma and Goto, 2001; Jack, 2004; Weigel and Meyerowitz, 1994) that captures and predicts floral whorls organ identity which is also verified at bud-burst and bloom through internode elongation (T_Long_).

Bud decompression permutation (T_Long_) (e.g. internode elongation) *in situ* (i.e. *in planta*) is crucial to both the SAM and the FM. It is minimal in the FM (a determinate growth organ system) and usually extensive in the SAM (an indeterminate growth organ system) (Benya and Windisch, 2007; Parcy et al. 2002).

The phenomenon of paleochronic reversion (i.e. an atavism) has been recognized fairly recently (Benya and Windisch, 2007). Goal of this research was to document, measure and chronicle any floral axial permutation (*in situ*) on paleochronically reverted floral specimens originating from multiple recombinants (*in planta*) of the species *Psophocarpus tetragonolobus* (L.) DC (fam. Fabaceae). Recombinants represented the two similar but significantly distinct environments where this reversion has been confirmed (Benya, 2012; Benya and Windisch, 2007). Analysis identified any significantly (SPSS, 2013) intense axial elongation activity at floral anatomic zones and/or regions.

## 2. Materials and methods

Data came from 70 paleochronically reverted floral specimens at a phylloid and/or phyllome ground state (Fig. 1 [right] and 2) originating from field-grown homeotic segregants (i.e. homozygous, recessive recombinants) (Benya, 1995) of the species *Psophocarpus tetragonolobus* (L.) DC (Fabaceae). Segregants were managed for sequential collection of floral specimens for purposes of timing analysis *in situ* or postharvest so that “reversion age” of floral specimens served as a timing mechanism. Definition of reversion age in days (RAD) is the time from initiation of reversion (day zero) on the recombinant, to the harvest date (and conservation) of any reverting floral specimen from that recombinant. It is the time, counting from the onset of reversion on the recombinant to the date when a floral specimen entered the laboratory in a post-harvest cluster (Benya and Windisch, 2007), in accord with similar procedures (Lohmann et al. 2010). Pre-whorls and whorls anatomic zones, both juxtaposed (Fig. 1) (Besnard et al. 2014) and with phyllotactic alteration (Fig. 2 and 3) (Pinon et al., 2013) (e.g. axil “permutatively distanced” [Benya, 2012]) were examined and characterized for decompression longitudinal (T_Long_), rotational (T_Rtn_) and/or latitudinal (T_Lat_) permutation. Changes were qualitatively recognized by location within any of two anatomic zones (the pedicel-bracts zone and the floral whorls zone) and of five morphologic regions (fields), sub-regions (sub-fields) and active structures therein. These included bracts (Bt), calyx (Cl), corolla (Crla), androecium (Andr), gynoecium (Gynec) and components therein (e.g. gynophore and/or cupule-like structure). Quantitative count of active sites per specimen then served to measure intensity and distribution of permutation.

**Figure 1.**
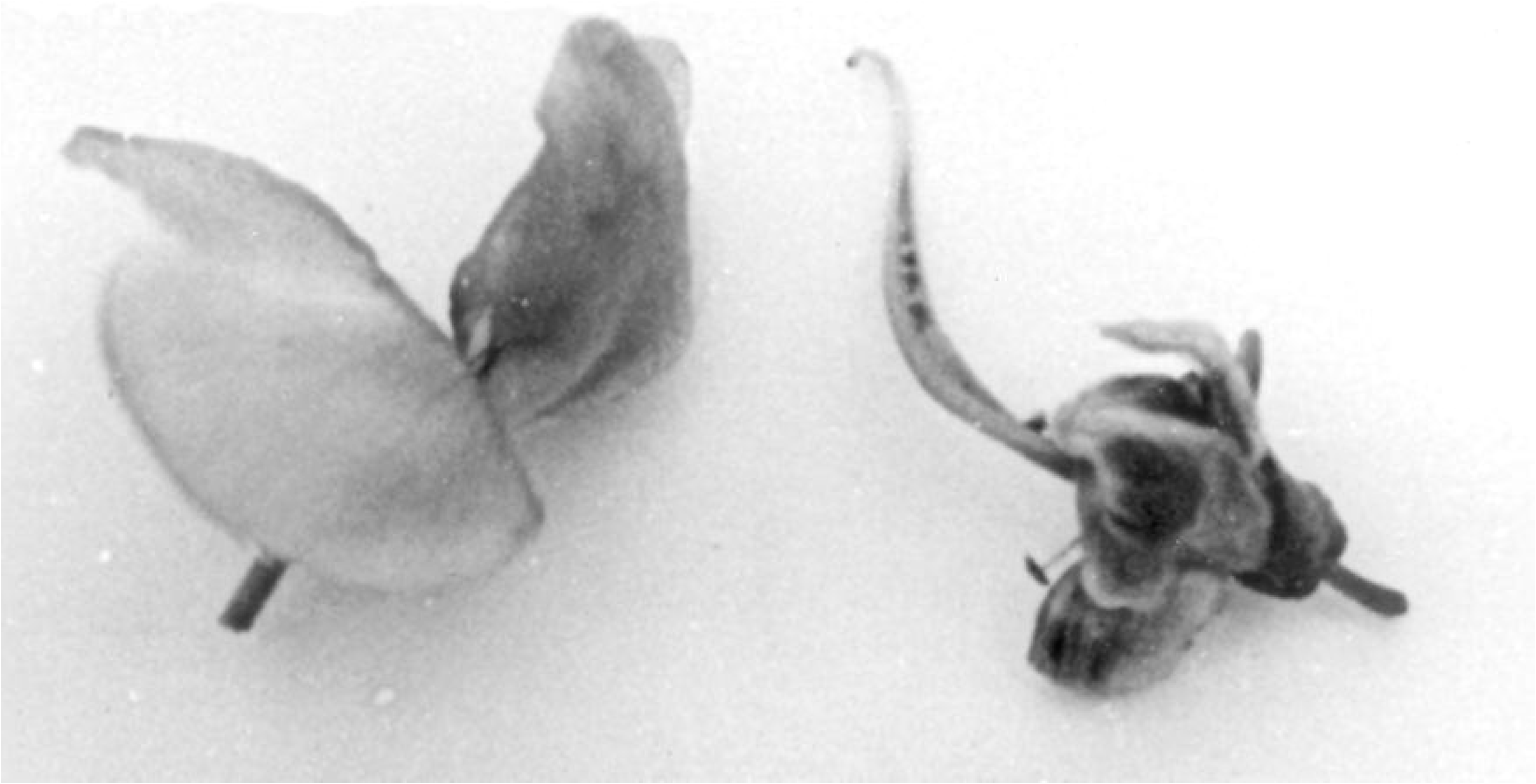
Flowers *Psophocarpus tetragonolobus*; normal (i.e. wild type; [left]) and reverted (i.e. a phylloid state, meristematically inactive, bracts and calyx regions juxtaposed; [right]).

**Figure 2.**
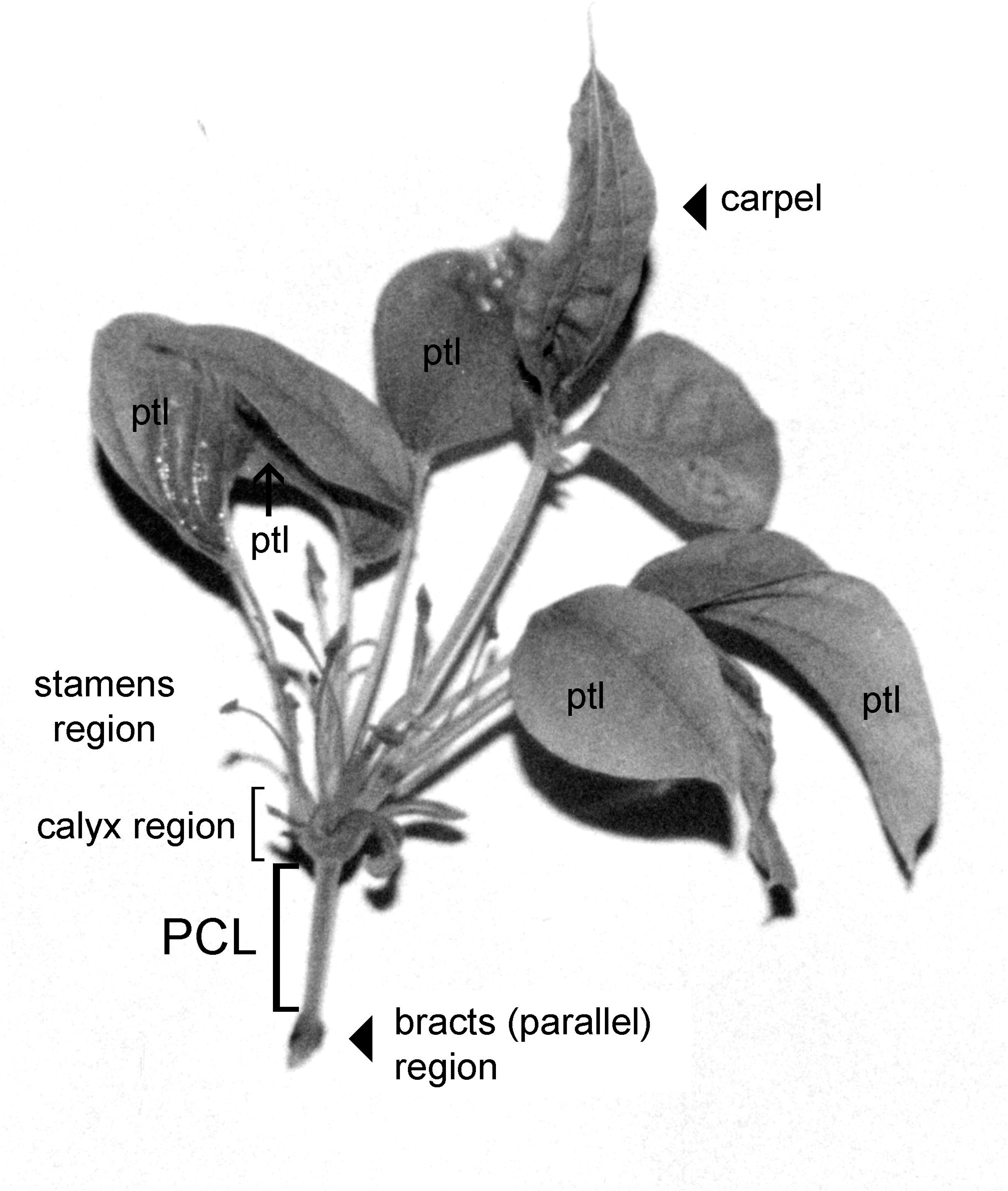
Reverted flower presenting beginning phyllome condition of whorls organs and permutation: carpel (top) petals (ptl) [sides] stamens (normal) [stamens region] calyx region [sepals] pericladial stalk (PCL) bracts (parallel) and calyx regions distanced about 1 cm from each other.

Specimens in laboratory were divided over 33 clusters containing one to four reps per cluster primarily oriented to maintaining specimens for purposes of physical measurement and determining any age and/or timing variables influencing activity. Treatments were conducted in individual glass test-tubes and/or plastic containers in plain or deionized water within simple-structured laboratories. Humidity and temperatures within the laboratories followed closely those of external environmental conditions during the winter and spring seasons in the semi-arid, tropical equatorial climates of Teresina, Piaui, (05°05’S; 42°49’W, alt. 64 m), from early August to late November (Gadelha de Lima, 1987) and late September to mid November in Russas, Ceará Brazil (04°55’S; 37°58’W, alt. 20 m).

### 2.1 Anatomic morphologic sequence

#### 2.1.1 Juxtaposition

Two floral anatomic zones (i.e. pre-whorl pedicel-bracts and whorls) presented six initial regions that served to begin clarifying results by anatomic location. Both zones are specific in their “normal” organs, the respective regions of those organs and the sequence(s) that define those regions. Specimens could have parallel bracts (Bt), juxtaposed to the whorls zone; calyx (Cl), corolla (Crla), androecium (Andr) and gynoecium (Gynec); juxtaposed, linearly compact (Fig. 1 [right]) reverted specimens. Categories of permutation were then based on those from published data (Benya and Windisch, 2007).

#### 2.1.2 Inter-zonal permutation

The possibility of phyllotactically altered specimens arose where floral axes were permutated presenting bracts that were physically “distanced” from the calyces by a pericladial stalk (PCL) resulting in inter-zonal longitudinal decompression (Fig. 3).

**Figure 3.**
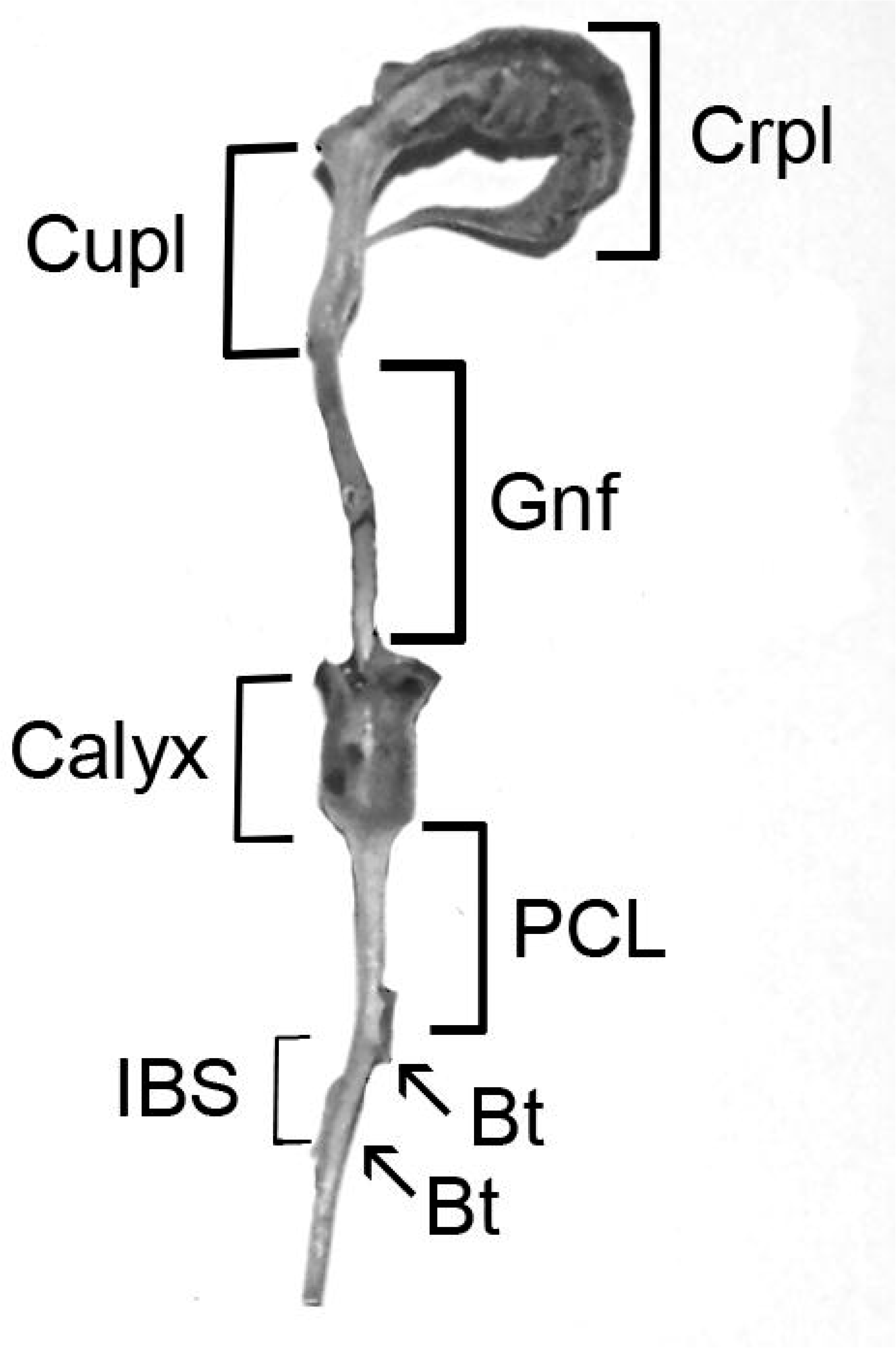
Reverted flower: Bracts (Bt) and bracts dislocation forming an “inter-bracts stem” (IBS) of about 4 mm (measured from locus center of one bract to locus center of second bract); Pericladial stalk (PCL) of about 8 mm distancing pedicel-bract zone from whorls zone; Calyx (petals and stamens removed for clarity); Gynophore (Gnf) of about 12 mm connecting to a Cupule-like structure (Cupl) of about 8 mm leading to a webbed carpel (Crpl) showing vascularization and initial spiraling (Benya and Windisch, 2007, Suppl. Material, photo amplified, cropped and adapted [photoshop] for this figure).

#### 2.1.3 Inter-regional permutation

A third scenario postulated decompression of specimens with bracts dislocation due to development of an inter-bracts stem (IBS) (Fig. 3) (Benya and Windisch, 2007).

Because of their recognition as defined floral whorls (Coen and Meyerowitz 1991; Parcy, et al. 1998; Schwarz-Sommer et al. 1990), each whorl is initially treated as a distinct region for possible permutation activity notwithstanding the actual permutation documented at each. Thus four additional categories of putative decompression might occur at the floral whorls zone; the calyx, corolla, androecium and the gynoecium with its component structures especially at the carpel (Fig. 4, 5, 6) and components therein. The point of conjunction of the androecium and the gynoecium could give rise to development of a gynophore (Gnf). In possible sequence with the gynophore a cupule-like structure (Cupl-Lk) might precede the carpel (Fig. 3). The structure is termed “cupule-like” because its homology is not yet confirmed. Thus six putative fields of permutation 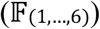 are hypothesized herein. Recognized site permutation that could not be measured exactly (e.g. partially emerged cupule-like or gynophore from a reverted “non-blooming” floral bud), received the measure of what emerged or a minimum measure of one mm (1 mm) for purposes of statistical analysis. Latitudinal site displacement (e.g. carpel cleft webbing) was recognized but not measured because of *in situ* impossibility of such measurement while still maintaining such specimens viable, as explained below.

**Figure 4.**
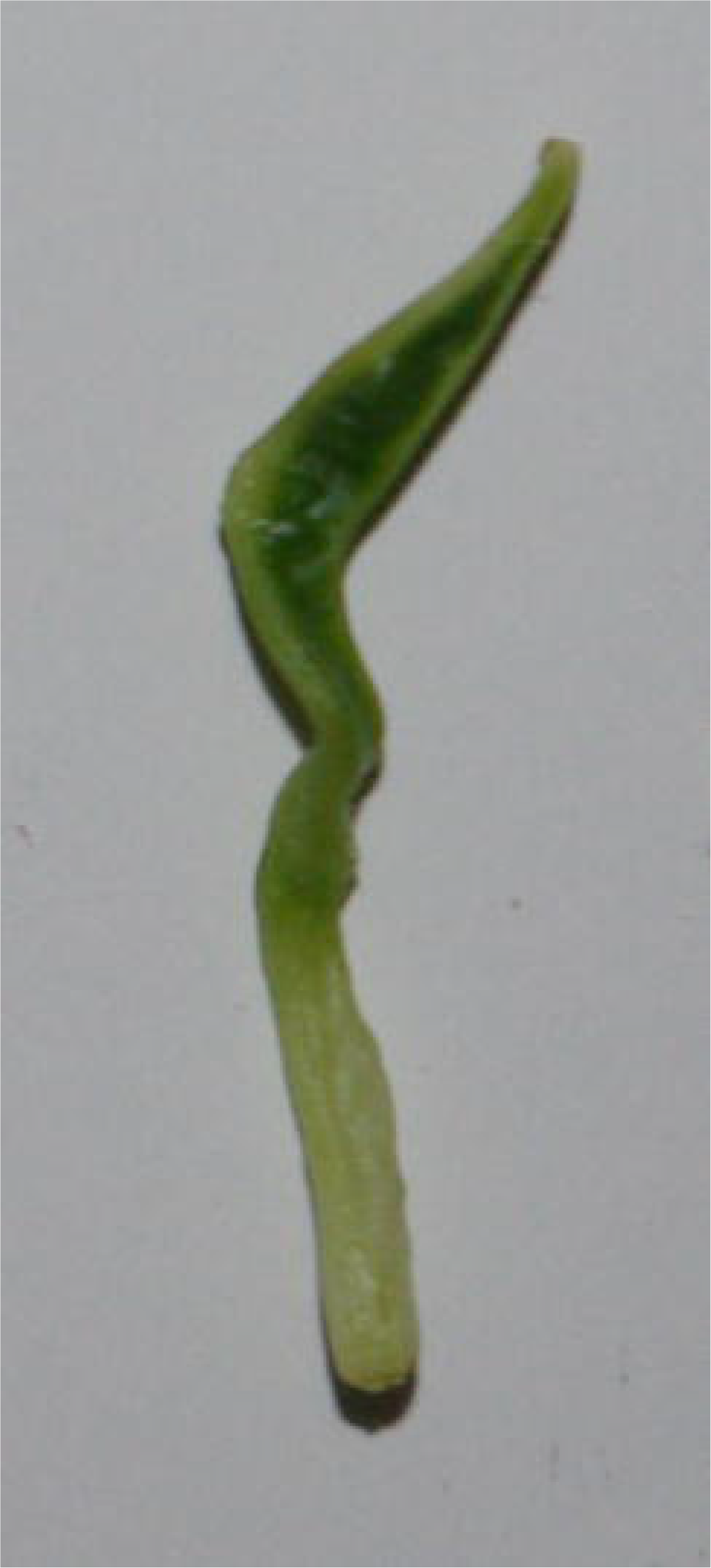
Webbing between carpel clefts is a first necessary step in permutation at the carpel.

**Figure 5.**
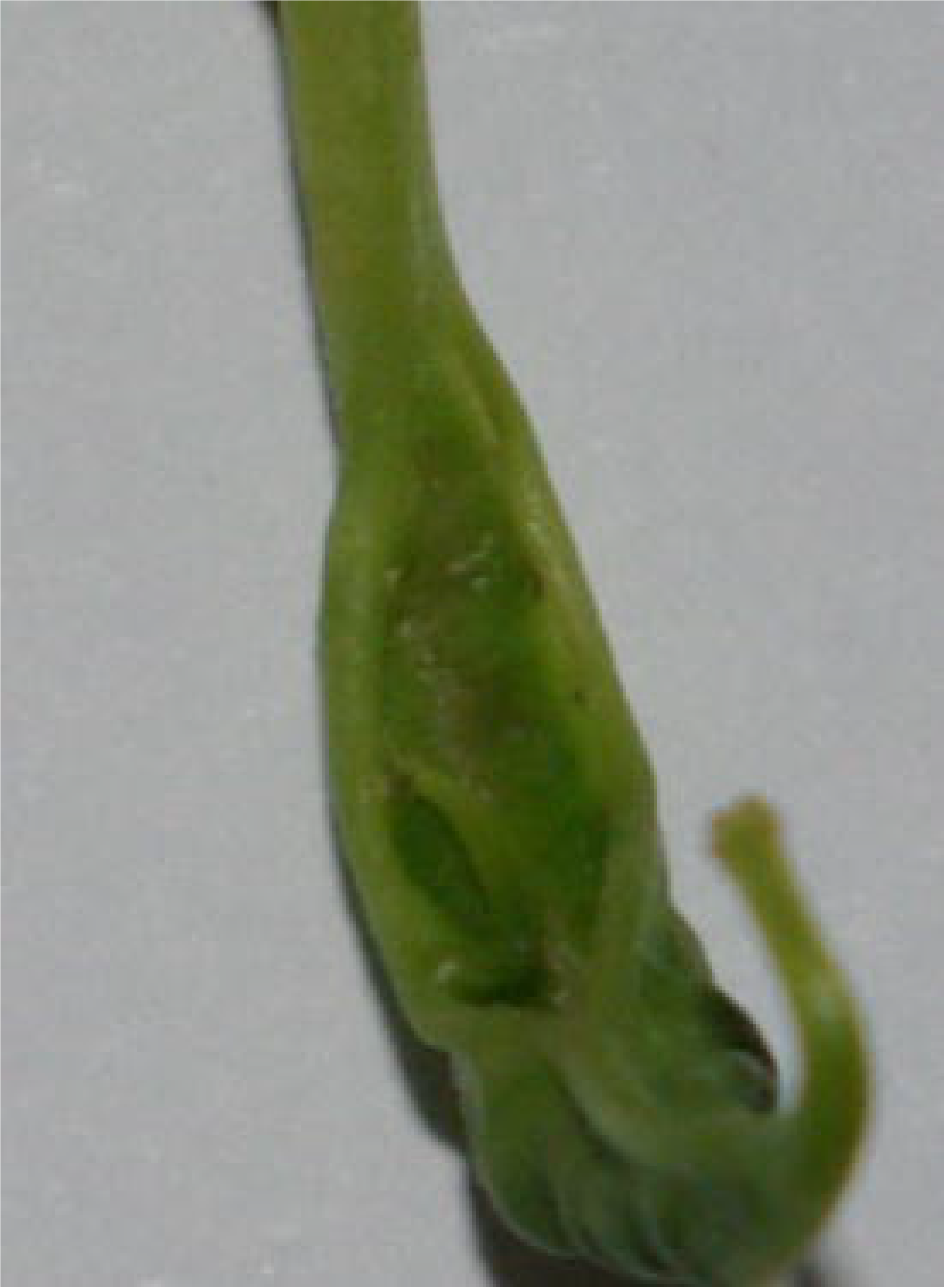
Vascularized carpel with initial diadnation of about 60%, acropetal along adaxial cleft.

**Figure 6.**
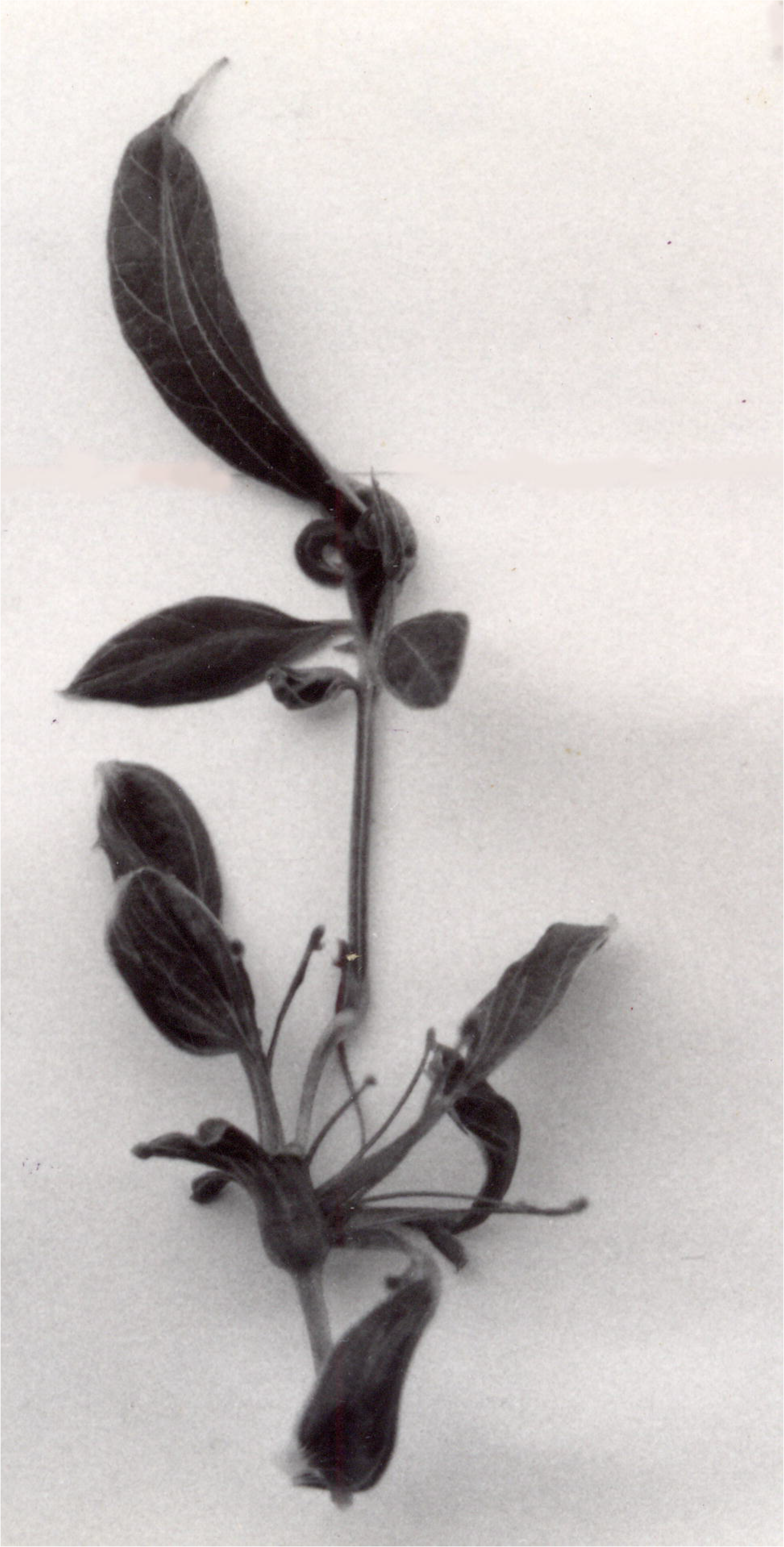
Reverted flower: Permutated carpel on extended cupule-like structure presenting the four necessary permutation steps leading to complete carpel foliation: 1. Webbing between carpel clefts; 2. Vascularized webbing 3. Carpel deadnation (acropetal along adaxial cleft); 4. Initial carpel foliation (putative ovules).

**Figure 7.**
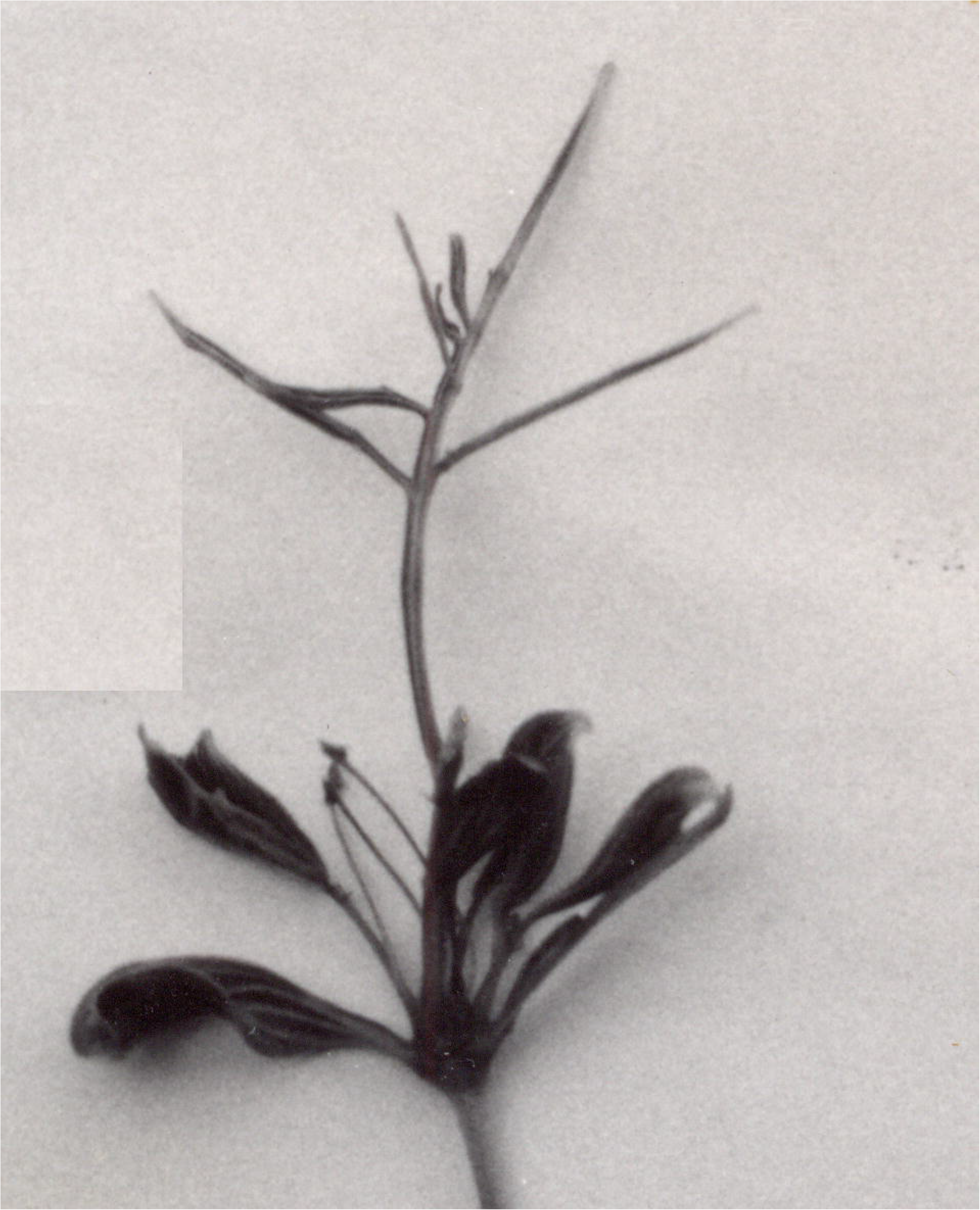
Reverted flower: Parallel carpel clefts, no webbing, definite diadnation, showing foliation presenting a “pinnate” carpel structure (top) [referential notation of photo removed for simplicity].

## 3. Results

A general transformation permutation (T_x_) arose as vascularization, diadnation, foliation (T_Phyld_) plus axial decompression (T_Axl_); as longitudinal (T_Long_), rotational spiraling (T_Rtn_) and/or lateral (T_Lat_) webbing function. This function varied in its diversity and distribution between the two anatomic zones (ANOVA, F_7, 62_ = 2.749, *p* = 0.015), among the six regions within those zones (χ^2^ = 467.732, *p* ≤ 0.005, df = 5) and its distribution within anatomic regions (ANOVA, F_31, 38_= 2.370, p ≤ 0.006). Permutation usually occurred *in situ* (*in planta*). However it could continue into laboratory. Remeasurement of a sample of 25 specimens in laboratory showed a total of 7.0 mm of PCL and/or IBS development among two specimens; one mm of PCL and six of IBS, statistically not significant (χ^2^ = 0.0144, NS, df = 1). Thus decompression in situ (immediate post-harvest) was taken as the overall measure of permutation in accord with similar procedures (Piao et al. 2015).

Loci of the bracts anatomically defined the bracts region which extended from zero (i.e. bracts parallel or normal) (Fig. 2) to 12 mm depending on dislocation and development of any IBS (Fig. 3). This occurred on 31 specimens. Inter-zonal decompression yielded a PCL which developed in lengths of one to 38 mm on 60 specimens. This was usually accompanied (but at times preceded or succeeded) by formation of an IBS since distinct genes determine the phenotypic presence of each structure as already reported (Benya and Windisch, 2007). One or both of these then constituted the “Axial active” (*n* = 63 Σ = 807.0 mm) measure of permutation activity on most specimens. PCL and IBS dislocations were distinct. Each could occur separately (i.e. three IBS and 32 PCL) on different specimens or concurrently on 28 specimens as did cupule-like structures while gynophore development occurred on nine specimens.

The sum of the lengths of the IBS and PCL (i.e. “Axial active” elongation) plus those of the gynophore and cupule-like structure formed the “Axial complete” measure (*n* = 65 Σ = 1115.0 mm) of “longitudinal decompression”. “Axial active” was the principal, significant (F_31_, 38 = 7.611, *p* < 0.000) component (72.38%) of “Axial complete” decompression whose length was more specifically constituted by PCL lengths (F_30, 29_ = 4.430, *p* < 0.000).

Linear axial displacement of a locus or loci in an established direction or directions (i.e. along the floral axis) was driven by axial expansion (AE). AE thus defined the principal demarcation components and all of the measured metric components (1115.0 mm) of longitudinal (T_Long_) permutation (128 of 274 sites) where each “demarcation event is a vector” (Green and Baxter 1987). A spiral (T_Rtn_) displacement of carpels of the gynoecium arose (*n* = 6 specimens), a topological dislocation vector function (rotational axial elongation). Webbing between carpel clefts (n = 39) is a lateral axial vector function (T_Lat_). Both T_Rtn_ and T_Lat_ are components of AE. However all three present signs of distinct vector functions. Their sum total equals axial complete expansion (T_Axl_); so that:

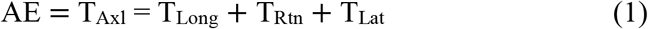

Regional decompression on two of the seven juxtaposed bracts-calyx specimens presented cupule-like structures. Thus the final ratio of confirmed axial decompression permutation to non-axial decompression specimens was 65:5.

Overall permutation activity was significantly more intense within the whorls zone (183 sites) than within the pre-whorls zone (91 sites) (*t* = 2.360, *p* = 0.021, df = 69). Research then focused on anatomic regions within both zones, the intensity and possible sequence(s) of decompression within and between regions, plus any distinctions in decompression between the two significantly different environments (Benya, 2012). “Axial complete” measure (AE = 1115 mm) of longitudinal floral decompression occurred over 2421 “reversion age in days” (RAD) at a mean value of 0.461 mm RAD^-1^ with a range of 0.003 to 3.563 mm RAD^-1^.

The difference between the 128 decompression sites and the 146 general permutation sites was not significant (χ^2^ = 1.1825, NS, df = 1). Timing of both decompression and general permutation functions at the gynoecium was significant across and within both environments (F_24_, 39 = 2.586, *p* ≤ 0.004). Cupula-like structuring was significant only in Russas (F_4, 11_ = 13.105, *p* < 0.000). Significance continued in the diversity of sites and events at and within the carpel (F_24, 39_ = 3.097, *p* ≤ 0.001). This included webbing between carpel clefts only at Teresina (F_19, 28_ = 2.530, *p* = 0.013), vascularization at both locations (F_24, 39_ = 2.765, *p* ≤ 0.002), then only at Teresina; diadnation and foliation, both at (F_19, 28_ = 2.395, *p* = 0.018), and putative ovule permutation (F_19, 28_ = 2.919, *p* ≤ 0.005), (also as vascularization, elongation, and/or foliation) in sequence and/or combination with some or all of these antecedent functions. This reflected the structural complexity of the carpel and the diversity of activity that could occur at that sub-region (Benya, 2012, Trigueros et al. 2009).

Parallel non-webbed carpel clefts at both environments preceded any permutation function at the carpel and probably preceded permutation at any other anatomic region. This is the “ground state” of the carpel. Permutative decompression of the carpel begins with webbing between carpel clefts (Fig. 4) and/or spiraling (Fig. 3). Both webbing and spiraling were virtually impossible to time through RAD in either environment as both were in situ functions and usually preceded *in planta* flower bloom thus impeding simple empirical verification. They were, however confirmed as “initiating functions” by means of dissection of pre-bloom flowers (i.e. flower buds) whose further permutation activity was thus eliminated because of dissection.

Amplification at the calyces (*n* = 8) (*t* = − 14.049, *p* < 0.000, df = 69), foliation of petals at the corolla (*n* = 2) (*t* = − 31.104, *p* < 0.000, df = 69) (Fig. 2), plus foliation of the anther (*n* = 2) (*t* = − 31.104, *p* < 0.000, df = 69) at the androecium all arose at low levels (T_Phyld_). Gynophore development was minimal over both environments (*n* = 9) (*t* = − 8.765, *p* < 0.000, df = 69). Cupule-like structure development followed a norm (*n* = 28) (*t* = −1.097, *p* = 0.276, NS, df = 69) commensurate with decompression at the IBS (*n* = 31) (*t* = − 0.119, *p* = 0.905, NS, df = 69). However it showed significant negative correlation (*r* = − 0.353; *p* ≤ 0.004) with RAD because it arose early in the decompression sequence.

Permutation as decompression and vascularization whose origin from the ground state carpel presenting a rigorous sequence of steps significantly grounded in the genetics of decompression and vascularization is already addressed at the phenotypic level (Benya and Windisch, 2007). However its manifestation (e.g. gene activation) is significantly influenced by weather (Benya, 1995) and climate (Benya, 2012). Thus the sequence is rigorous but not invariable especially because of alleles of genes governing aspects of axial decompression phenotype in a dominant:recessive Mendelian scenario (Benya and Windisch, 2007).

Homozygous recessive recombinants (extremely rare) could affect that sequence of steps, excluding entire steps in the sequence (i.e. webbing and/or vascularization of the carpel [a dominant phenotype]) in a multi-recessive homozygous recombinant, thus giving rise to diadnate carpels showing no webbing and no vascularization and presenting “pinnate” carpel form (Fig 7 [top]). The lateral axial function (T_Lat_) was attenuated or completely inactive. However the prevalence of dominant alleles plus their manifestation allowed reasonable deduction of sequence pertaining to preceding phenotypes according to established precedence (Benya and Windisch, 2007). These could terminate at any step as: un-webbed to webbed, then perhaps to vascularized, then at times to diadnation, then sometimes to foliation (Fig. 6) (Table 1).

**Table 1:**
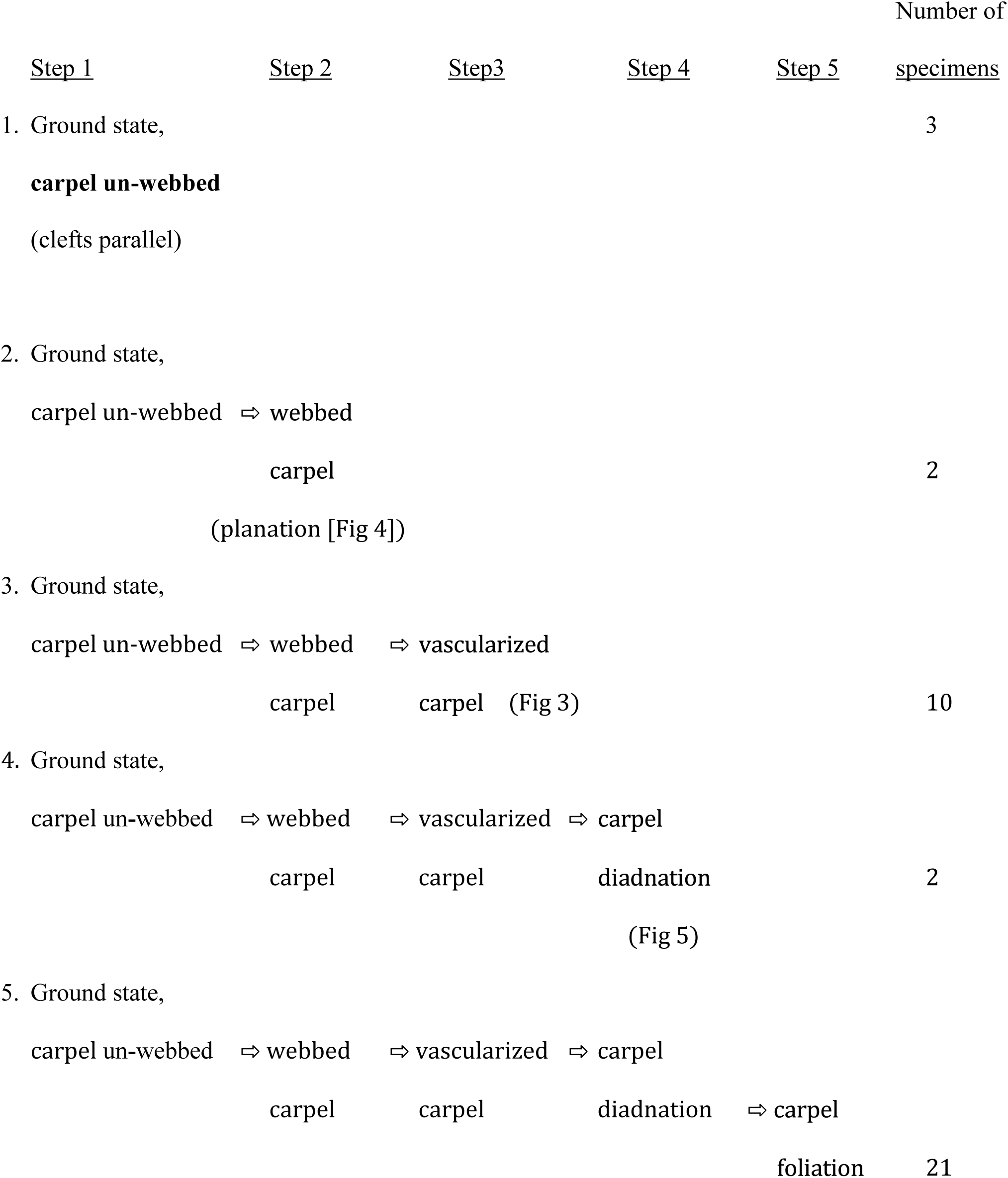
Putative “necessary structural permutation and developmental steps” within the carpel, begin with ground state, un-webbed, parallel carpel clefts and progress in steps, each of which can be terminal OR continuous to the next step.

Two general sequences of permutation activity are manifest in this data. The first sequence involves metric intensity of activity between anatomic zones, regions and subregions beginning at the bract-calyx juncture. Overall intensity of site activity (n = 274) followed a significant quadratic regression on axial complete decompression (1115 mm) (quadratic r^2^ = 0.301, F_2, 67_ = 14.4 3 5, *p* < 0.000) from bracts to whorls inclusive. That regression continued for the four floral whorls themselves (*n* = 183 sites) (quadratic r^2^ = 0.230, F_2, 67_ = 10.024, *p* < 0.000), into the fourth whorl gynoecium (*n* = 171 sites) (quadratic r^2^ = 0.197, F_2, 67_ = 8.213, *p* ≤ 0.001) and total carpel structures (*n* = 134) (quadratic r^2^ = 0.101, F_2, 67_ = 3.7 7 3, *p* = 0.028).

The T_Lat_ function of webbing (n=39) (linear r^2^ = 0.070, F_1, 68_ = 5.103, *p* = 0.027), vascularization (*n* = 37) (linear r^2^ = 0.073, F_1, 68_ = 5.326, *p* = 0.024), and diadnation (*n* = 27) (linear r^2^ = 0.110, F_1, 68_ = 8.399, *p* ≤ 0.005) presented a sequence of significant linearly varying intensity, while carpel foliation (*n* = 25) (quadratic r^2^ = 0.104, F_2, 67_ = 3.898, *p* = 0.025) and internal carpel foliar number (*n* = 61) (quadratic r^2^ = 0.103, F_2, 67_ = 3.85 9, *p* = 0.026) presented significant quadratic regression on the axial complete value. Spiraling, not a necessary function of the carpel sequence, occurred at a non-significant level (six specimens) and only at Russas.

A second sequence involved timing. This placed webbing of the carpel (usually in Teresina) and/or spiraling of the carpel (usually in Russas), neither being strictly elongation functions, as initiatory or co-initiatory events closely followed by minimal but early calyx amplification on the pre-bloom flower. Diadnation and foliation of the carpel could then follow webbing in as little as 24 hours. Where carpel vascularization occurred, it usually preceded diadnation and foliation of the carpel. Vascularization is governed by a dominant allele (*VASCARP*) (Benya and Windisch, 2007). However activation of that allele depended on weather and climatic conditions (Benya, 1995, 2012). Gynophore and/or cupule-like structural formation might follow webbing.

IBS and/or PCL elongation (i.e. Axial active = 807 mm) in relation to RAD was significant, (ANOVA F_24, 39_ = 2.083, *p* = 0.020). It could be almost initiatory. However its significantly broad physically spatial distribution as a component of Axial complete length; 807 of 1115 mm (ANOVA F*31, 38* = 7.611, *p* < 0.000), rank it among the most time consuming of events in relation to RAD (r = 0.195, *p* = 0.122, *n* = 64).

As in the case of site analysis in relation to axial metric decompression (Axial complete = 1115 mm), regression analysis revealed dynamics of site establishment in relation to RAD of specimens. Overall intensity of site activity; phylloid and/or AE (*n* = 274 sites) showed significant response to RAD (ANOVA F_24, 39_ = 2.285, *p* = 0.011) but no significant tendencies (i.e. linear, quadratic, etc.) (Suppl. Table 1). However whorls site activity (n = 183) (r^2^ = 0.100, F_2, 61_ = 3.374, *p* = 0.041), activity at the gynoecium (*n* = 171) (r^2^ = 0.096, F_2, 61_ = 3.256, *p* = 0.045), structural development internal to the carpel (*n* = 134) (r^2^ = 0.150, F_2, 61_ = 5.392, *p* ≤ 0.007) and even carpel spiraling (*n* = 6 sites) (r^2^ = 0. 214, F_2,61_ = 8.282, *p* ≤ 0.001) all presented significant quadratic response to RAD.

The plethora of dynamic activity in response to RAD beginning at the whorls zone, showed progressive intensity therein continuing into the gynoecium region and into the carpel sub-region (plus loci therein). It presented chronologic significance that was quadratic in all cases even including spiraling of the carpel (Suppl. Table 1). That plethora of activity seems to capture a dynamic whose basis lies in the progressively intensive genetic governance already identified for the whorls and carpel (Álvarez-Buylla et al., 2010; Ashman and Majetic, 2006; Coen and Meyerowitz, 1991; Prunet et al., 2008; Schwarz-Sommer et al., 1990; Weigel and Meyerowitz, 1994) and even implying genetic aspects yet to be recognized.

## 4. Discussion

A phylloid ground state and/or various degrees of phyllome organ formation (Pelaz et al., 2000; Weigel and Meyerowitz, 1994) characterized all 70 experimental specimens. Floral meristem cancellation (Benya and Windisch, 2007), anticipated and was essential to that phylloid state. After that, most specimens entered into a permutation phase of transformation (T_x_) that could include organ foliation (T_Phyld_) (Fig. 2) and/or a decompression function (T_Axl_) of the floral axis, itself constituted by axial elongation (T_Long_), rotational (T_Rtn_) and/or lateral (T_Lat_) dynamic of permutation (Okabe 2011, 2015). The permutation function was significant *in situ* (*in planta*) but could extend to postharvest. By deduction, it usually began in the carpel as webbing between carpel clefts (T_Lat_) and/or rotational spiraling (T_Rtn_). This was usually prior to flower bloom, thus specific (*in vivo*) timing and measurement of these two events was impossible. However significant negative linear correlation of carpel spiraling and near significant correlation of carpel webbing (r = − 0.416, *p* ≤ 0.001, *r* = − 0.221, *p* = 0.079 respectively) with RAD support this conclusion. Significant linear correlation of whorls zone active sites (n=183) with axial complete permutation (r = 0.480, *p* < 0.000) verify both the dynamic of whorls site activity and the intensity of that activity as informed by pre-reversion site canalization at the whorls zone.

Bracts regions usually entered a morphologically active phase of elongation (Bargmann et al., 2013; Benya and Windisch, 2007; Besnard et al., 2014; Pinon et al., 2013). These were the most striking in their manifestations of the permutation elongation phase as IBS and/or PCL. However decompression as gynophore and/or cupule-like structure formation contributed to overall axial elongation. Rare yet at times solitary presence of PCL, IBS and cupule-like structures indicated that a distinct vector governs each of these decompression events. Those distinctions coincide with Mendelian proportions indicating dominant:recessive genetic functions underlying IBS and PCl presence or absence (Benya, 2012; Benya and Windisch, 2007). Significant negative linear correlation of cupule-like structures, number (*n* = 28, r = − 0.353, *p* ≤ 0.004) and length (*n* = 211, r = − 0.659, *p* ≤ 0.001), with RAD but their positive correlation with Axial complete measure, respectively (r = 0.381, *p* ≤ 0.001) and (r = 0.509, *p* ≤ 0.007), verifies their initiation early in the elongation process. It further supports the premise that their origin is through distinct genetic governance. Specific governance at the gynophore cannot be determined from this data.

Bracts (with any IBS therein) plus any PCL show continuity to the calyces presenting a succession of distinct fields and a sub-field 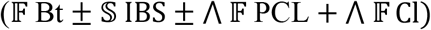 representing temporal and physically spatial activity. The PCL is a biophysical field 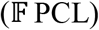 whose varying morphologic length, serves to distance the Cl field 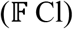 and Bt field 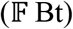 from each other. The anatomic regions 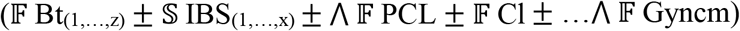 constitute a permutated floral axis (Axl), beginning at Bt regions and extending to the carpel (Crpl) of the gynoecium inclusive. It is a biophysical continuum of function, but not a AE structural continuum. The physical sequence of fields 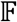 Bt_(1,…,z)_ …, 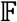 Cl show AE continuity, but no AE occurred at the corolla or androecium where permutation was limited to the phylloid (T_Phyld_) function. AE continuity was interrupted. It arose again at 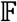 Gyncm 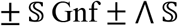 Cupl-Lk. Thus the anatomic region 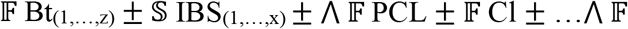 Gyncm is a continuum of function. It is a dynamic longitudinal linear axial vector space 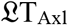 but not a continuous AE structural space.

Besides distancing bracts from calyces, a PCL also distanced the entirety of the floral pre-whorls pedicel-bracts anatomic zone from the whorls anatomic zone. Results indicate definite regional homeostatic canalization associated with paleochronic floral reversion (Benya and Windisch, 2007). Canalization continued to be manifest as floral permutation up to and including axial elongation at these anatomic zones and their respective organs regions (Okamuro et al., 1993).

Lack of any significant correlation between RAD and permutation activity at the calyces, corollas and androecium reflects their robust phenotypes arising from canalization of pre-reversion organ identities (Debat and David, 2001; Okamuro et al. 1993) and resulting stability at these regions. Intensity of activity diminished between the calyx (eight specimens) and gynophore (nine specimens) to a minimum at the corolla and androecium (two specimens each). It then increased from nine at the gynophore to 28 specimens with a cupule-like structure and then to the 63 specimens with a total of 134 carpel permutation sites. Cubic regression thus reflected the inversely varying robusticity of organ identity (due to pre-reversion canalization) with permutation function from pre-whorls into whorls floral sites. Presence but lack of any significant relation of the spiraling function (*n* = 6) with overall permutation site activity (*n* = 274) reflects the distinction between whorls structuring (and any whorls structuring function) and the spiraling function (Okabe, 2011). The sequence was completed by the invariably consistent negative linear correlations of overall site activity, whorls site activity (Suppl. Table 1) plus regions therein including gynoecium and carpel spiraling, webbing and vascularization with RAD. It verified the intensity of decompression activity early in the permutation phase as lability of established floral morphologic compression following paleochronic reversion. That lability diminished over time.

Juxtaposition of floral anatomic zones and organs regions is the phyllotactic norm for this and most other angiosperm species. During elongation, floral organs maintain their specified identities at definite positions of their respective loci (Benya and Windisch, 2007). However expansion of anatomic organ regions, by means of PCL, IBS, gynophore, etc. can augment organ regional longitudinal dimensions and even change locus orientation and fields.

## 5. Conclusion

Sexually reproductive flowers can revert (transmutation) from the determinate growth reproductive state to a non-reproductive phylloid state. Reverted flowers can then enter a permutation phase where physical spacing of organ regions occurs along and within the floral axis. Distinct biophysical functions affect that permutation phase.

Spiraling function in the SAM can be captured with mathematical precision (Okabe, 2011) while overall SAM genesis can be captured in a simple model (Young, 1978). That model can be expanded in the FM to a distinct ABC(DE) model of homeotic gene function for floral whorls genesis. Cancellation of the dynamic of the ABC(DE) model leads to a floral ground state. A permutation function can then arise. The permutation functions documented here manifest significant and variable presence in a significantly specific but variable timing sequence as reverted flowers demonstrate a return to indeterminate growth through AE (axial expansion) in this “Axial permutation” model. This model contains both axial decompression and foliation components

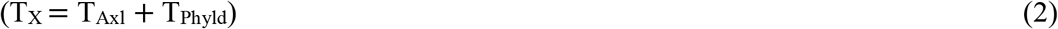

the prior of which is composed of axial longitudinal (T_Long_), spiral (T_Rtn_) and latitudinal (T_Lat_) components so that:

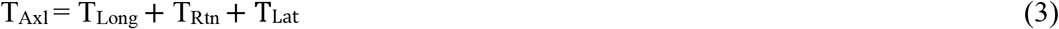

Where

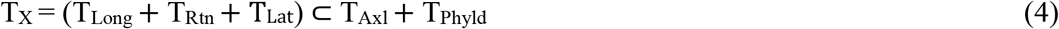

Metrics of T_Long_ are documented in this data. Presence of T_Rtn_ and T_Lat_ are qualitatively recognized but quantitative aspects of each are not addressed herein. Thus:

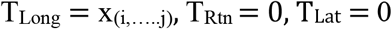

where:

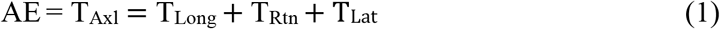

so that:

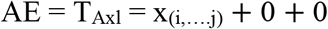

Thus since T_Rtn_ and T_Lat_ are effectively “zero” in this data, therefore:

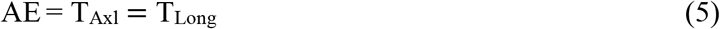

Variability of the ABC(DE) model is due to variable activity of homeotic genes. Variability of this “Axial permutation model” is also due to homeotic genes (Benya and Windisch, 2007) but their activation is significantly correlated with climatic and weather factors (Benya 2012). Resulting anatomic sequence of permutation activity then runs from the bracts (Bt) region to the carpel inclusive with components therein. The formula:

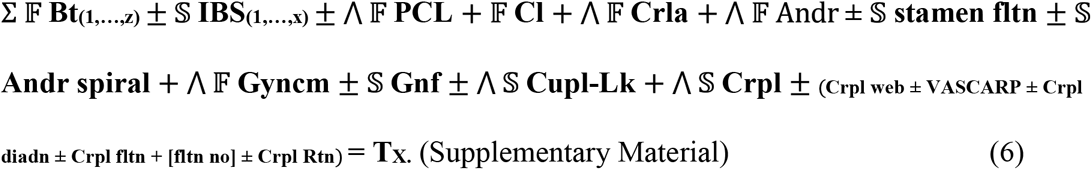

summarizes an anatomic sequence of permutation transformation (T_x_) in its phylloid (T_hyld_) and floral axial decompression (T_Axl_) aspects. This includes elongation (T_Long_), latitudinal (T_Lat_) and/or rotational (T_Rtn_) functions. The principal components of the longitudinal axial vector space (T_Long_) within this model (T_Axl_) are captured by the formula:

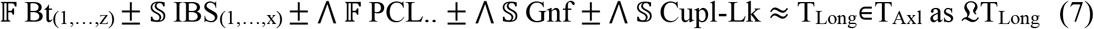

The early lability of floral form following paleochronic reversion hearkens to the unusually high labile floral phyllotaxis in ancestral angiosperms (Endress and Doyle, 2007). The presence of both linear, spiral, and latitudinal functions in this model and their distinct responses to permutation and timing variables (Suppl. Table 1) may be unique for paleochronically reverted flowers. The question of their simultaneous or sequential presence is not resolved by this data.

Ancestral reference further supports the distinction between decompression and spiral functions documented by this data. The spiraling function then gives rise to the question of its origin; primitive or derived (Endress and Doyle, 2007). However, distinction of longitudinal, latitudinal and topologic functions seems quite clear with the suggestion that it may well be primitive. Research in fact has been such that “. developmental studies have focused on vegetative rather than floral phyllotaxis because vegetative shoot apices are technically more tractable than floral apices in model plants.” (Endress and Doyle, 2007; Okabe, 2011, 2015). Combining both foci (i.e. SAM and FM) may be quite possible through the use of paleochronically reverted organisms.

Biophysical functions affect the permutative phase at the anatomic and morphologic regions studied here. A continuum might extend to further morphologic fields generated on flowers of species whose bract numbers increase in multiples beyond the dual-bract flower structure addressed herein. Theoretically that continuum could be extended longitudinally in segments (i.e. linear spaces) of varying lengths defined by each bract in a flower of multi-bract species (e.g. *Euphorbia pulcherrima, Cornus florida, Quercus* sp.) whenever the master “*srs*” recessive allele (Benya and Windisch, 2007), homozygous and activated, is accompanied by the necessary “reversion dependent genes” (Benya, 2012; Benya and Windisch, 2007). Each bract would thus define a specific linear field (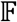 Bt_(1…z)_) with possible accompanying sub-fields of IBS (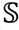 IBS_(1…x)_) with any PCL (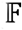 PCL) as part of the overall formula from bracts to carpel 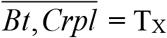 in anatomic sequence constituting a statistically dynamic yet mathematical vector space 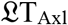 whose presence on living specimens of extant species offers a unique tool for research.

## Acknowledgements

Saint John’s College, Landivar, Belize City, Belize, (T. Thompson, professor) provided an introduction to material. G.R. Lovell (Griffin, Gerogia, USA), W. Denny (Beltsville, Maryland, USA) USDA-ARS, T.N. Khan, Dept. Agr. Western Australia and H.P.N. Gunasena, U. Peradeniya, Sri Lanka provided seed. A.C. Machin assisted with seed importation. Escola Agrícola Santo Afonso Rodriguez, (J. Moura Carvalho, E.M. Moreira, J. Bulfoni and I. Govoni) and Escola Técnica Soinho provided facilities for experimentation. The “Universidade do Vale do Rio dos Sinos” (UNISINOS) and “Laboratório de Histologia” (Ana Leal-Zanchet, coordinator) furnished facilities for analysis. A. DePaula, J. M. daSilva, E.O. Alves, J. deFreitas, C.G. deOliveira, F. Gil and G. & H. Galik helped with technical work and analysis. P G. Windisch, M.C. Moura Carvalho, C. Radz, S.J.V. Benya and T. H. Oliveir assisted with manuscript preparation.

